# Joint estimation of relatedness coefficients and allele frequencies from ancient samples

**DOI:** 10.1101/103903

**Authors:** Christoph Theunert, Fernando Racimo, Slatkin Montgomery

**Affiliations:** Department of Integrative Biology, University of California, Berkeley; Department of Evolutionary Genetics, Max-Planck Institute for Evolutionary Anthropology, Leipzig, Germany; New York Genome Center, New York, NY 10013

## Abstract

We develop and test a method to address whether DNA samples sequenced from a group of fossil hominin bone or teeth fragments originate from the same individual or from closely related individuals. Our method assumes low amounts of retrievable DNA, significant levels of sequencing error and contamination from one or more present-day humans. We develop and implement a maximum likelihood method that estimates levels of contamination, sequencing error rates and pairwise relatedness coefficients in a set of individuals. We assume there is no reference panel for the ancient population to provide allele and haplotype frequencies. Our approach makes use of single nucleotide polymorphisms and does not make assumptions about the underlying demographic model. By artificially mating individual genomes from the 1000 Genomes Project, we determine the numbers of individuals at a given genomic coverage that are required to detect different levels of genetic relatedness with confidence.

## Introduction

Over the past few years the amount of ancient DNA (aDNA) recovered from fossilized bones, teeth and hair has grown rapidly [12], [16], [10]. Despite significant advances in sequencing technology, laboratory practices and computational methods, problems still arise because of low amounts of endogenous nuclear DNA, short degraded fragments, contamination from present-day humans and sequencing error. Nevertheless, data from ancient remains are a precious source of information, providing insights about the history of humans and their closest relatives that are unavailable from any other source. DNA from several ancient individuals found in the same location are especially important because they can provide clues about relatedness within groups. This information is valuable for downstream analyses which make assumptions about relatedness among individuals.

In sexually reproducing species, the coefficient of relatedness (*r*) is twice the probability that two sites sampled at random from autosomes (one from each individual) are identical by descent (IBD). With that definition, *r* = 1 for two samples from the same individual or from monozygotic twins, *r* = 1*/*2 for first-degree relatives (parents and offspring or full siblings), *r* = 1*/*4 for second-degree relatives (aunt or uncle and nephew or niece, half siblings, grandparent and grandchild, or double first cousins), etc.

Information about the genetic relatedness between individuals is of significance in the fields of forensic sciences, agriculture, human genetics and ecological sciences. A variety of approaches have been developed to infer relatedness, each suited to specific types of data. For a comprehensive review on statistical methods and available approaches see [20] and [17]. The general concept underlying relatedness analyses is that of IBD, but this quantity cannot be observed directly in data. Instead allelic states at a particular locus are used to make inferences about IBD and relatedness. When good quality, high-coverage genomes from individuals are available, inferring relatedness is relatively easy and many methods have been developed (for example [11], [13], [19], [4], [2], [7]). However, for ancient DNA, the quality and amount of data are often sufficiently limited that existing methods cannot be applied.

There have been some attempts to deal with the problems posed by ancient DNA. For example, Vohr et al. developed an approach to identify whether two DNA samples with extremely low coverage originate from the same or different individuals [18]. The authors introduced a likelihood method that uses information from single nucleotide polymorphisms (SNPs) and patterns of linkage disequilibrium. However, this method relies on a reference panel of phased haplotypes from the same population in order to infer allele and haplotype frequencies. This method can be used for human fossils that are sufficiently recent that a present-day population can be used as a reference panel. However, for older human fossils and for Neanderthals and Denisovans, no reference panels are yet available.

In another recent study, Martin and Slatkin [9] presented a method to infer close genetic relatedness using low-coverage next generation sequencing (NGS) samples from ancient individuals. They did not assume a reference panel is available and they account for contamination from modern humans and sequencing error. Their method investigates the overlap of pairwise genetic distance distributions calculated under certain realistic scenarios to identify the relatedness between pairs of individuals.

In this study we extend the work of [9] by using a maximum likelihood framework applied to each polymorphic site and determine whether this approach provides improved accuracy. In addition to inferring relatedness, our method provides estimates of allele frequencies and contamination levels for each sample. By artificially mating individual sequences from the publicly available 1000 Genomes Project we determine the number of individuals at a given genomic coverage that are required to distinguish different levels of genetic relatedness.

## Methods

### Model notation

The relatedness *r* of two individuals is twice the probability of identity by descent of two chromosomes chosen at random. Individuals are denoted by *i, j* = 1, 2*,…, N* and sites are denoted by *k* = 1, 2*,…, L*. We further assume the sequencing error rate *e* is the same for every site *k* in every sequence. *e* is the probability that a site is misread during the sequencing, if it is misread at a site that is actually monomorphic then it creates a false SNP, but if it is misread at a site that is actually polymorphic then it is misread as the alternative allele. The contamination rate for an ancient individual *i* is *C*_*i*_. *C*_*i*_ is the probability that a randomly chosen read from an individual *i* is derived from a present day human. The average contamination rate over all sequenced ancient individuals is denoted 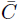. We use only sites that are polymorphic in the contaminant panel and we will assume that we observe only ancestral or derived (non-chimpanzee) alleles at every site, thereby ignoring triallelic sites.

Furthermore, let *f*_*k*_ be the derived allele frequency (*daf*) at site *k* in the putative contaminating population (e.g. modern humans). The observed *daf* at site *k* in the ancient samples is *q*_*k*_ and is a weighted average of the endogenous and contaminating allele frequencies:

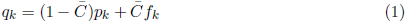
where *p*_*k*_ is the endogenous *daf* at site *k* in the ancient samples (unobservable because the alleles sequenced at a site might either be endogenous or from the contaminating population). We use 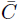 because it is in principle impossible to determine which of the reads at a given site comes from the contaminating population. Therefore, our best estimate of *p*_*k*_ is:

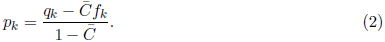

Summarizing, the model input parameters are the allelic (ancestral/derived) states at each site from each of the ancient reads. The observed parameter is *q*_*k*_. *f*_*k*_ is the only parameter used from a contaminating reference dataset. The more individuals that are available in the contaminating reference dataset the closer these values approach true population frequencies resulting in more accurate parameter estimates. A parameter that cannot be directly observed from the ancient data is *p*_*k*_, but it is calculated at each step based on *q*_*k*_, *f*_*k*_ and 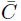. The parameters that we will aim to estimate are the relatedness coefficient for each pair of individuals *r*_*i,j*_, the contamination rate for each individual *C*_*i*_ and the overall sequencing error rate *e*.

For a pair of individuals *i* and *j* with relatedness *r*_*i,j*_, there are three sets of parameters that need to be modeled.

1. Endogenous frequencies - the probabilities of allelic configurations 11,10,01,00 in the ancient DNA (1 being derived, 0 being ancestral):

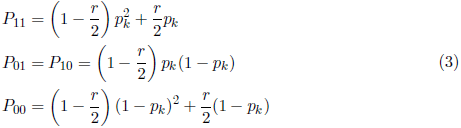
2. Contaminated frequencies - the probabilities of allelic configurations in the contaminated sample:

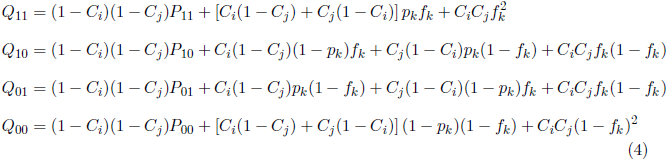
3. Sequenced frequencies - the probabilities of allelic configurations in the sequences themselves, allowing for sequencing error:

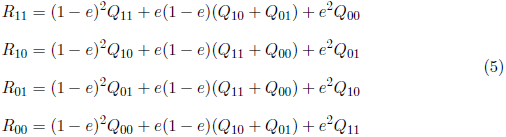

### Parameter estimation

Assume a dataset of *N* ancient individuals and *L* aligned sites. For each pair of individuals *i* and *j* (out of *N*(*N* − 1)*/*2 total pairs) the log likelihood is calculated as 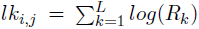. The log likelihood for the entire dataset is then the sum over all log likelihoods *lk*_*i,j*_ for all pairs of individuals. We refer to this approach as the “complete method”.

Overall the number of values that need to be estimated are *N*(*N* − 1)*/*2 parameters for the relatedness coefficients *r*_*i,j*_, N parameters for contamination rates *C*_*i*_ and one parameter for the sequencing error rate *e*. A method to maximize the log-likelihood of these input parameters is implemented in C++ using the non-linear optimization routine *L-BFGS* from the dlib C++ library [5]. The software package we generated will be made available online. Lower and upper bounds for the parameters *r*, *C* and *e* are set to [0.001, 0.9999] [0.0, 0.25] and [0.001, 0.25] respectively.

As mentioned in the results, a slightly different procedure of using only a subset of all available individuals to calculate the likelihood for the entire dataset was tested. In this case *n < N* individuals are used to calculate the overall likelihood as the sum over *n*(*n* − 1)*/*2 likelihoods *lk*_*i,j*_. For example, if one is only interested in a certain pair of individuals *i* and *j*, then *n* = 2 and only one *r*_*i,j*_ needs to be estimated. However, *q*_*k*_ at site *k* is still estimated using all N individuals. Depending on the actual value of *n* this approach may result in faster computation times. We refer to this approach as the “subset method”.

We simulated 50 independent datasets for each combination of *N*, *L* and *r* and separately performed the parameter estimation for each of them. Therefore, the final estimates of *r* are given as an average, and the accuracy of our method is evaluated by the root-mean-square error (*rmse*) and the mean absolute error (*mae*). When used together, *rmse* and *mae* can characterize the errors in a set of forecasts. The magnitude of the difference between them is informative about the amount of variance in the individual errors in the sample.

### Simulations

For the initial evaluation of the model we generated sets of *2N* sequences of length *L* sites. For each sequence, alleles at each position *k* were either derived (1) with probability *P*_*k*_ or ancestral (0) with probability (1 − *P*_*k*_), where *P*_*k*_ was randomly drawn from *U*[0, 1]. In order to generate contaminated reads from our simulated genotypes we adopt a method used in [14]. For each simulated individual *i*, the number of derived and ancestral fragments at a particular site follows a binomial distribution that depends on the true ancient genotype, the sequencing error rate and the contamination rate *C*_*i*_ (see equations 3-6 in [14]). Contamination rate *C*_*i*_ for individual *i* was randomly drawn from a uniform distribution between 2% and 25% separately for each simulation (i.e. in each simulation individuals have different rates of contamination *C*_*i*_). Sequencing error rate *e* was set to 0.001 throughout all simulations. To systematically study the behavior of our method we assume one read per individual at each simulated genomic position. We further assume *f*_*k*_ for each site from a putative reference panel to be randomly drawn from *U*[0, 1].

Furthermore, we simulated a scenario where reads are only available from a random subset of individuals (out of a total of N) at each genomic site. Supplementary tables 3 and 4 summarize the results.

To ensure the simulation method mentioned above does not introduce any biases we carried out simulations where we artificially mated unrelated European (EUR) sequences from the 1000 Genomes Project Phase 3 [1]. A similar approach was introduced in [9]. We focused on phased genomes and extracted all biallelic polymorphic sites from single chromosomes from EUR individuals. Contamination from a putative contaminant panel was implemented in the same way as described before. We restricted our analyses to SNPs that passed the basic 1000 Genomes Project filtering criteria and for which ancestral allele information was available. The ancestral states were determined by using information from the inferred human-chimpanzee ancestor at each site. We filtered sites with a Map20 < 1 (Duke uniqueness tracks of 20bp) and we removed deletions and insertions. The method behaves exactly the same for both datasets (simulated sequences and sequences from the 1000 Genomes Project).

In both cases, we performed artificial meioses of pairs of individuals for single chromosomes. The recombination rate was assumed to be uniform along the genome and set to be 1.310^−8^ per bp per generation [6],[12]. We implemented a minimal number of one recombination event per chromosome per generation. Relatedness among individuals was then simulated by artificially mating them with other individuals to produce offspring.

To investigate the effect of different types of genomic sites, we analyzed each dataset (not the present-day reference panel) filtered for 1) fixed and polymorphic sites 2) only polymorphic sites and 3) polymorphic sites that were either changed to being fixed or remained polymorphic after allowing for contamination and sequencing error. This way, we could study the effect of different classes of sites on the accuracy of our estimates.

### Simulated pedigrees

Three different pedigrees (denoted *f1*, *f2* and *f3*) were generated with the mating method described above:

- f1: father + mother = child
- f2: father + mother = child1; child1 + X0 = child2
- f3: father + mother = child1; father + mother = child2; child1 + X1 = child3; child2 + X2 = child4

where X0, X1 and X2 represent unrelated partner individuals. Throughout the manuscript each dataset is further represented by an additional number which denotes the absence (.0) or presence (.1) of contamination and sequencing error (e.g. *f2.1* is the second pedigree *f2* with contamination and error). In each dataset all remaining individuals were kept unrelated. The very last individual in each dataset is a direct copy of another individual before contamination and error. This allows us to test the method for different degrees of relatedness: *r* = 0.5 in f1; *r* = 0.5 and *r* = 0.25 in f2; *r* = 0.5, *r* = 0.25 and *r* = 0.125 in f3; and *r* = 1.0 in all three.

## Results

### Accuracy when *C*_*i*_ = 0 and *e* = 0

First we studied the accuracy of our method to identify genetic relatedness simulated in pedigree *f1.0* in the absence of contamination and sequencing error by using the subset approach with *n* = 2.

In figure 1 each subfigure of boxplots represents a different combination of *N* individuals (rows) and *L* sites (columns) and shows the distribution of *r* for a pair of related individuals over 50 independent datasets (see supplementary figure SF1 for more details and error values). Note that we refer to the true simulated relatedness coefficient as *r*_*s*_, the point estimates of it as *r* and the estimated average over 50 independent datasets as 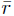.

**Figure 1:**
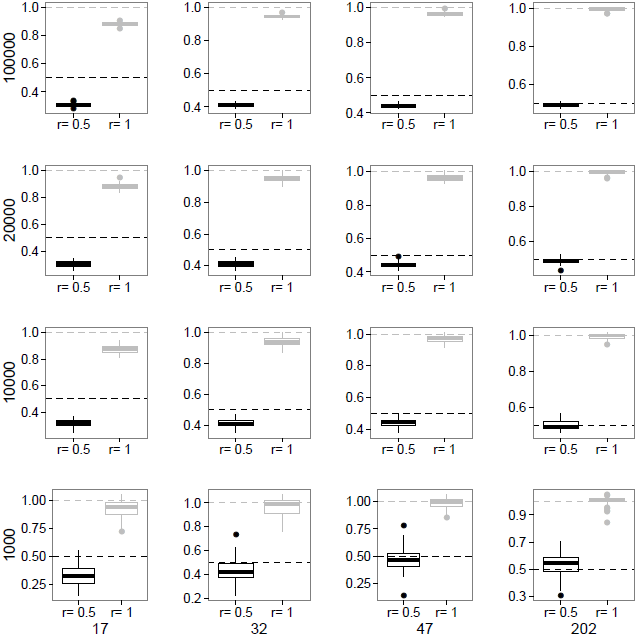
Each panel represents a different combination of *N* (columns) and *L* (rows) and shows a boxplot for the estimates of simulated relatedness of *r*_*s*_ = 1.0 and *r*_*s*_ = 0.5 over 50 independent datasets for pedigree *f1.0*. Dashed horizontal lines denote the simulated values of *r*_*s*_.

For example in the upper right corner we simulated reads for 202 diploid individuals and 100,000 overlapping polymorphic sites. For the two individuals that result from the same individual (*r*_*s*_ = 1.0), estimates are 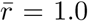 with rmse = 0.01 and mae = 0.01. In the same dataset for a pair of parent-offspring individuals (*r*_*s*_ = 0.5), 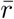 is 0.49 with rmse = 0.01 and mae = 0.01.

The variation of the parameter estimates is given in more detail in supplementary figure SF2 showing that the range in estimates for this dataset is rather small (for *r*_*s*_ = 1.0: r = [0.98,1.01]; for *r*_*s*_ = 0.5: r = [0.47,0.49]). We note here that values of *r >* 1 are possible as the final step of the parameter inference is *r*_*i,j*_ = *r*_*i,j*_*/*1 − (*mean*(*C*_*i*_*,C*_*j*_)). In the majority of cases the method underestimates the value of *r*_*s*_. As expected, the fewer overlapping sites and individuals that are available, the more the estimates deviate from the true value of *r*_*s*_ and the higher the error estimates become. For example for N = 17 and L = 1,000, 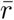 is 0.94 with rmse = 0.11, mae = 0.09 (for *r*_*s*_ = 1.0); and 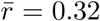 with rmse = 0.2 and mae = 0.18 (for *r*_*s*_ = 0.5). It is worth mentioning that the distribution of estimates for different *r*_*s*_ do not overlap with each other in any of the datasets.

Supplementary figure SF3 shows the comparison of estimated and simulated contamination rates for each related individual with the rmse shown in the legend of each graph (note that the true *C*_*i*_ = 0). The method overestimates the contamination rates, but the majority of *C*_*i*_ is estimated to be *<* 0.05 when the number of individuals and sites increase.

The accuracy of the method to identify relatedness coefficients from pedigree *f2.0* is presented in figure 2. Again, with reads from 202 individuals and *L >* 1, 000 the results are highly accurate and simulated *r*_*s*_ = 0.5 as well as second degree relatedness *r*_*s*_ = 0.25 are estimated to be 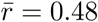 and 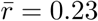 respectively (see supplementary figure SF4 for more details and error values).

**Figure 2:**
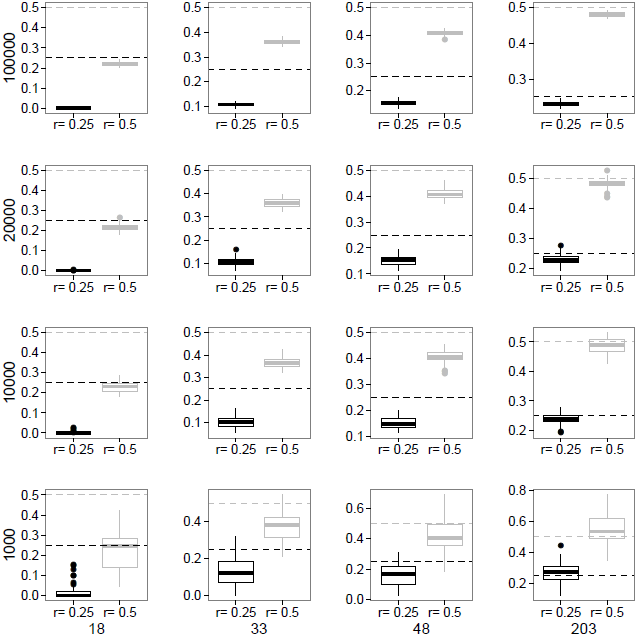
Each panel represents a different combination of *N* (columns) and *L* (rows) and shows a boxplot for the estimates of simulated relatedness of *r*_*s*_ = 0.5 and *r*_*s*_ = 0.25 over 50 independent datasets for pedigree *f2.0*. Dashed horizontal lines denote the simulated values of *r*_*s*_.

As expected, the smaller N and L, the less accurate results become, e.g. 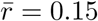 and error estimates around 0.10 for *r*_*s*_ = 0.25 when N = 48 and L = 10,000. Furthermore, with N = 18 the method does not pick up the signal of *r*_*s*_ = 0.25 anymore. Although for more distantly related individuals parameter inference may be less accurate, distributions of estimates do not overlap and so provide valuable information about differences in relatedness (see supplementary figures SF5 and SF6).

Identifying a relatedness of *r*_*s*_ = 0.125 from dataset *f3.0* is even more difficult. Shown in supplementary figures SF7, SF8 and SF9 are estimates for *r*_*s*_ = 0.125. It can be seen that only with reads from 205 individuals results are rather accurate at 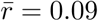 and errors of around 0.03.

### Accuracy when C > 0 and e > 0

Under a more realistic scenario, contamination from modern humans and sequencing error may create bias. Therefore we tested the method on simulated datasets that are affected by these factors. As described before, each *C*_*i*_ is drawn from *U* [0.02, 0.25] and *e* is set to 0.001 for all datasets.

Summarizing the information for pedigree *f1.1* from figures 3, 4, SF10 and SF11 the observations are similar to what we reported before but *C*_*i*_ and *e* affect the accuracy of the results. The method is still able to identify the same or related individuals while the amount of available data has a more pronounced effect on the accuracy. Estimates are less accurate than in the absence of *C*_*i*_ and *e*. However, when comparing the results for *r*_*s*_ = 1.0 and *r*_*s*_ = 0.5 from the same dataset, the distributions of estimates for *L >* 1, 000 do not overlap. This does provide valuable information (see figures SF10, SF11). For example, for 32 individuals and 10,000 sites, estimates of the relatedness coefficient range between [0.68, 1.08] when *r*_*s*_ = 1.0 and [0.2, 0.45] when *r*_*s*_ = 0.5. With *N >* 200 and *L >* 1.000 estimates of *r* and *C*_*i*_ are highly accurate with small error values.

**Figure 3:**
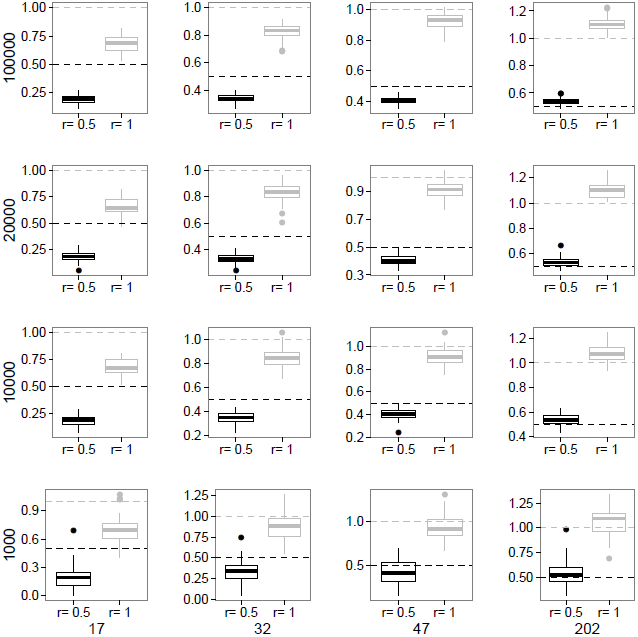
This figure presents results for estimates of *r*_*s*_ = 1.0 and *r*_*s*_ = 0.5 as in figure 1, but with *C*_*i*_ being between 2% and 25% and sequencing error set to 0.001 as in pedigree *f1.1*.

**Figure 4:**
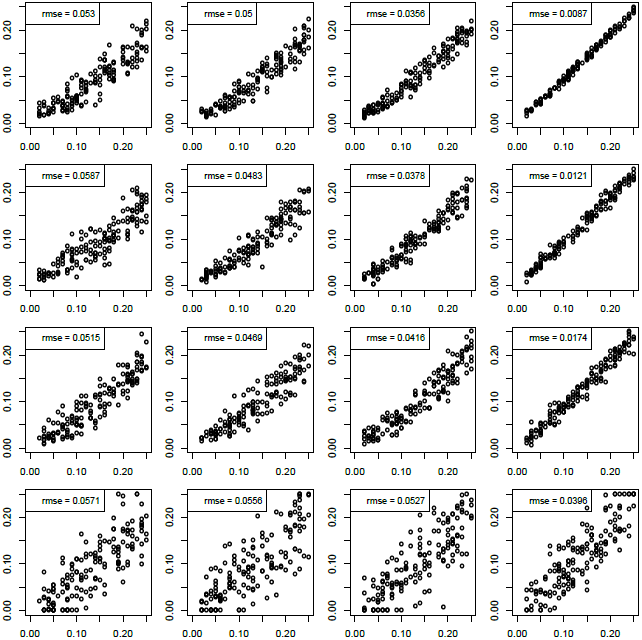
Each panel represents a different combination of *N* (columns) and *L* (rows) and shows an x-y plot for estimated (x axis) and simulated (y axis) contamination rates for pedigree *f1.1*. *C*_*i*_ was simulated to be between 2% and 25%.

The more distant the genetic relatedness between two individuals the more data are needed to identify it. Figure 5 shows results for pedigree *f2.1*. Again, note that with 203 individuals and ≥ 10.000 sites, *r*_*s*_ of 0.25 and *C*_*i*_ are accurately inferred (see supplementary figures SF12, SF13 and SF14 for more details and error values). The same is true for pedigree *f3.1* as seen in supplementary figures SF15, SF16 and SF17. The likelihood landscape under the presence and absence of *C*_*i*_ and *e* is shown in supplementary figure SF30.

**Figure 5:**
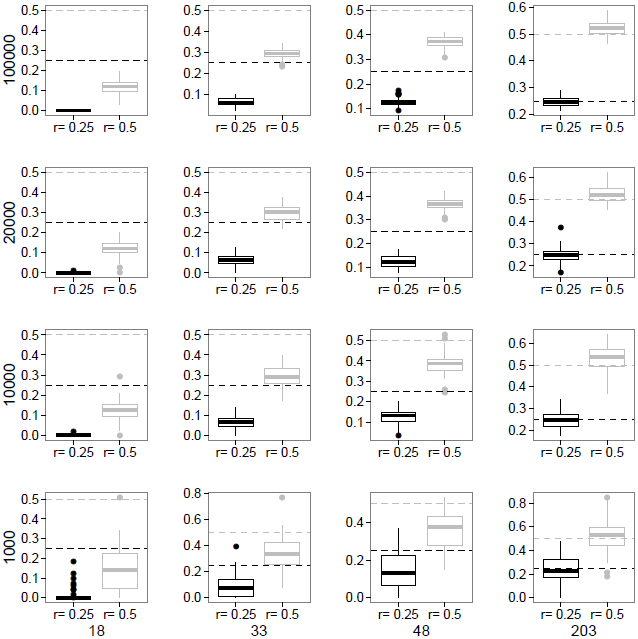
Estimates of *r*_*s*_ = 0.5 and *r*_*s*_ = 0.25 as in figure 2, but with *C*_*i*_ being between 2% and 25% and sequencing error set to 0.001 as in pedigree *f2.1*.

In conclusion, the proposed method can accurately infer the degree of genetic relatedness even in the presence of contamination and sequencing error. However, 1st, 2nd and 3rd degree relatedness require more data to be identified than when identifying DNA sequences that originate from the same individual. For example, for a *r*_*s*_ = 1.0 and N = 32, 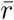 is still 0.84. In this case the distributions of estimates in figure SF11 show that the values do not drop below r = 0.7 in the majority of the cases (for *L >* 1, 000). Our method tends to underestimate the parameters without an overlap between the distributions of estimates for *r*_*s*_ = 1.0, *r*_*s*_ = 0.5 and *r*_*s*_ = 0.25. This fact supports the validity of results for *r*_*s*_ = 1.0 even more, as it seems unlikely that an estimated value of r = 0.8 is seen when the DNA sequences do not originate from the same individual. Even though the individual contamination rates are slightly overestimated, the final estimates of *r* are not heavily affected by this.

## Discussion

In this study, we present a method to infer the relatedness coefficients from aDNA samples sequenced from a group of fossil hominin bone or teeth fragments. Our method accounts for sequencing error and for contamination from present-day humans. By artificially mating simulated sequences as well as sequences from the 1000 Genomes Project, we determine how many overlapping reads and how many individuals are required to obtain estimates of relatedness coefficients with confidence. The likelihood model we developed for this purpose differs from existing methods in that we directly model the (hidden) ancient derived allele frequencies and do not require a reference panel for the ancient population.

In our simulations, we assumed that each polymorphic site is sequenced in every individual. With that assumption, the number of overlapping sites is a parameter under our control. The actual number of overlapping sites when there is low coverage sequence data is a random variable whose distribution depends on the sequencing method used. For shotgun sequencing, the simplest assumption is that the number of times a polymorphic site is sequenced is a Poisson distributed random variable with the mean equal to the coverage level, *λ*_*i*_ for individual *i*. The probability that the site is sequenced at least once in individuals *i* and *j* is (1 − *e*^*λ_i_*^)(1 − *e*^*λ_j_*^). For example if *λ*_*i*_ = *λ*_*j*_ = 0.1 (i. e. 0.1X coverage in both individuals), the probability that a site is covered by at least one read is roughly 0.009. Therefore if there are 3 ∗ 10^6^ polymorphic sites, there would be roughly 27,000 overlapping sites in two individuals. Different sets of sites would overlap in different pairs of individuals. Hence the expected number of samples that contribute to estimates of allele frequencies at each site in a sample of N individuals is 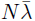 where 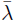 is the average coverage level. In the supplementary section (see supplementary tables 3 and 4 and supplementary figures SF18 - SF29) we allowed for this possibility in simulations by assuming that fully overlapping sites in all individuals are not available. As expected the accuracy of the method decreases when compared to using all individuals. However, the more individuals in total are available in a dataset, the higher the accuracy even when only using read information from a random subset (e.g. 5 or 10 individuals) of them at each genomic site. An alternative to shotgun sequencing is genomic capture ([8] and [15]). With a capture method that targets sites known to be polymorphic in the same or a closely related population, the probability that two sites are sequenced in two individuals depends on a number of factors, including the closeness of the population or populations used for ascertainment and the complexity of the genomic library. However, the success of targeted capture methods can be quite high. For example, Castellano et al. [3] used exome capture on two Neanderthal samples. In the El Sidron sample which had 0.2% endogenous DNA, 92.8% of targeted sites were covered at least once. In the Vindija 33.15 sample which had 0.5% endogenous DNA, 98.8% of the targeted sites were covered. Therefore, if exome or SNP capture methods are used there is a good chance of high levels of overlap in different individuals.

In our analysis we assume that the sampled (ancient) population is in Hardy-Weinberg equilibrium. That assumption allows us to derive the genotype frequencies from allele frequencies. If a population is made up of inbred individuals, then our method would not yield accurate results.

We do not make any inferences about the time of separation of the contaminating (present-day) population from the sampled (ancient) population. If the contaminating population is closely related to the sampled population, then allele frequencies in the two populations will be similar and the estimated allele frequencies in the ancient sample will depend only weakly on the contamination rate. The estimate of the contamination rate would not be accurate but the error in estimating that rate would not strongly affect estimates of relatedness coefficients. If the contaminating population has quite different allele frequencies, the estimates of contamination rate will be more accurate.

Finally, admixture from the ancient (e.g. Neanderthal) population into the contaminating (e.g. modern human) population will not affect our method. Admixture will make some of the contaminating allele frequencies slightly more similar to Neanderthals than they would be in the absence of admixture, which should not affect the estimates of contamination rates or relatedness coefficients.

**Table 1:** Estimation results are shown for a pair of simulated parent-offspring individuals when using the complete approach (n = N individuals) and the subset approach (n = 2 individuals). Numbers in the second and third column are as follows: 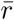 rmse of 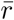; rmse of*C*_*i*_. 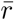 is given as an average over 50 independent runs. Although when using n = N, r is estimated for all possible pairs of individuals, only one is shown for the pair of interest.

| Dataset | n = N | n = 2 |
| --- | --- | --- |
| N=17, L=10.000, $C_i=0$, $e=0$ | 0.30, 0.21; 0.056 | 0.31, 0.19; 0.037 |
| N=32, L=20.000, $C_i=0$, $e=0$ | 0.40, 0.10; 0.02 | 0.41, 0.09; 0.019 |
| N=17, L=10.000, $C_i > 0$, $e > 0$ | 0.17, 0.33; 0.058 | 0.19, 0.32; 0.06 |
| N=32, L=20.000, $C_i > 0$, $e > 0$ | 0.31, 0.20; 0.047 | 0.35, 0.16; 0.056 |

## Acknowledgments

The authors gratefully acknowledge the help of Mike Martin. This work was supported in part by the Max Planck Society (as a salary for C.T.) and in part by a US National Institutes of Health grant (R01-GM40282 to M.S. and F.R.).

## Supplementary Material

Supplementary figures are available online at www….

Moreover, we compared the subset approach to the complete approach (see see ‘parameter estimation’ in Methods section). For example, for a complete set of N = 17 there are (17 ∗ (16)/2) = 136 different *r*_*i,j*_ to be estimated, one for each pair of individuals. When only interested in a subset of n = 2 individuals, there is only one *r*_*i,j*_ that needs to be estimated, greatly reducing the computational time. Table 1 shows results for two examples of 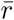 for a certain pair of parent-offspring individuals. We do not observe striking differences in the estimates and error values between the two approaches. However, estimates for *C*_*i*_ under pedigree *f1.1* are slightly better when using n = N as the allelic configurations of all individuals contribute to the overall likelihood. These differences in *C*_*i*_ account for the slight differences in estimates of r.

We further analyzed whether different types of genomic sites affect the results of the estimations. We tested the method by filtering the data for 1) fixed and polymorphic sites 2) only polymorphic sites and 3) polymorphic sites that were either changed to being fixed or remained polymorphic after biasing the data with contamination and sequencing error.

**Table 2:** Estimation results are shown for a pair of simulated parent-offspring individuals when using polymorphic and fixed sites (1) or only polymorphic sites (2). Numbers in the second and third column are as follows: *r*_*i,j*_, rmse of *r*_*i,j*_; rmse of*C*_*i*_. *r*_*i,j*_ is given as an average over 50 independent runs.

| Dataset | Class 1 | Class 2 |
| --- | --- | --- |
| N=17, L=10.000, $C_i > 0$, $e > 0$ | 0.15, 0.36; 0.10 | 0.19, 0.32; 0.059 |
| N=32, L=10.000, $C_i > 0$, $e > 0$ | 0.33, 0.18; 0.064 | 0.35, 0.16; 0.046 |
| N=47, L=10.000, $C_i > 0$, $e > 0$ | 0.41, 0.10; 0.039 | 0.40, 0.11; 0.040 |

**Table 3:** Subsampling approach: estimates of simulated relatedness coefficients (and rmse) of *r*_*s*_ = 1.0 and *r*_*s*_ = 0.5 over 50 independent datasets for pedigrees *f1.0* and *f1.1* when using reads from only 5 randomly selected individuals (out of N) per genomic site.

|  |  |  | N=5 of 17 | N=5 of 32 | N=5 of 47 |
| --- | --- | --- | --- | --- | --- |
| f1.0 | $r_s = 1.0$ | L=10.000 | 0.89, 0.15 | 0.97, 0.14 | 0.98, 0.17 |
|  |  | L=20.000 | 0.87, 0.15 | 1.02, 0.11 | 1.0, 0.09 |
| | $r_s = 0.5$ | L=10.000 | 0.31, 0.23 | 0.41, 0.21 | 0.42, 0.28 |
|  |  | L=20.000 | 0.31, 0.20 | 0.43, 0.14 | 0.48, 0.18 |
| f1.1 | $r_s = 1.0$ | L=10.000 | 0.71, 0.38 | 0.87, 0.26 | 1.01, 0.27 |
|  |  | L=20.000 | 0.70, 0.34 | 0.87, 0.23 | 0.96, 0.27 |
| | $r_s = 0.5$ | L=10.000 | 0.19, 0.32 | 0.36, 0.27 | 0.40, 0.36 |
|  |  | L=20.000 | 0.22, 0.30 | 0.35, 0.25 | 0.40, 0.23 |

We refer to these different classes of sites as 1, 2 and 3. As can be seen from an example in table 2, especially for datasets with fewer individuals and sites, using only sites from class 2 does slightly improve the estimates of contamination rates and, therefore, the overall estimates of *r*. In summary, accuracy for the different site classes is as follows: 1 *>* 3 *>* 2. We therefore apply the method to polymorphic sites only.

The results shown so far are based on an implementation of the method where *p_k_* is calculated as 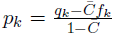. We also implemented an approach where each *p_k_* is a parameter that is estimated by the optimization algorithm and not calculated as before. This adds L parameters to the optimization procedure. This approach also strongly increases the computational time of the method. Obtained results (not shown) are surprisingly similar to the original method but because of the increase in runtime and lack of improved accuracy this technique is less useful in practice.

**Table 4:**
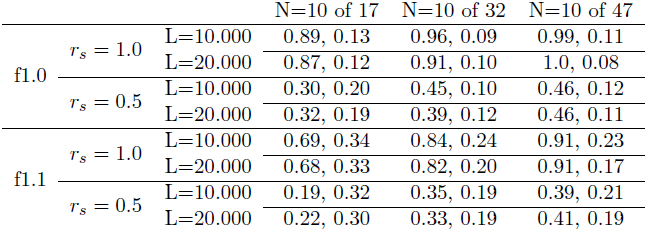
Subsampling approach: estimates of simulated relatedness coefficients (and rmse) of *r_s_* = 1.0 and *r_s_* = 0.5 over 50 independent datasets for pedigree *f1.0* and *f1.1* when using reads from only 10 randomly selected individuals (out of N) per genomic site.

---

**Figure SF1:**
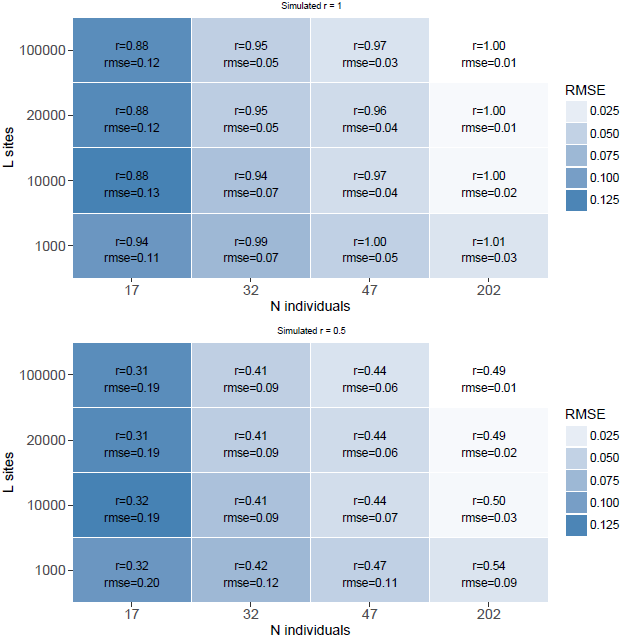
Each colored box represents a different combination of *N* and *L* and shows the estimated relatedness coefficient *r* (and *rmse*) for a pair of related individuals as an average over 50 independent datasets for pedigree *f1.0*. In each box we show results for simulated *r*_*s*_ = 1.0 (top panel) and *r*_*s*_ = 0.5 (bottom panel). *C*_*i*_ = 0, *e* = 0. The color intensity represents the magnitude of the *rmse* with darker color referring to a higher *rmse*.

**Figure SF2:**
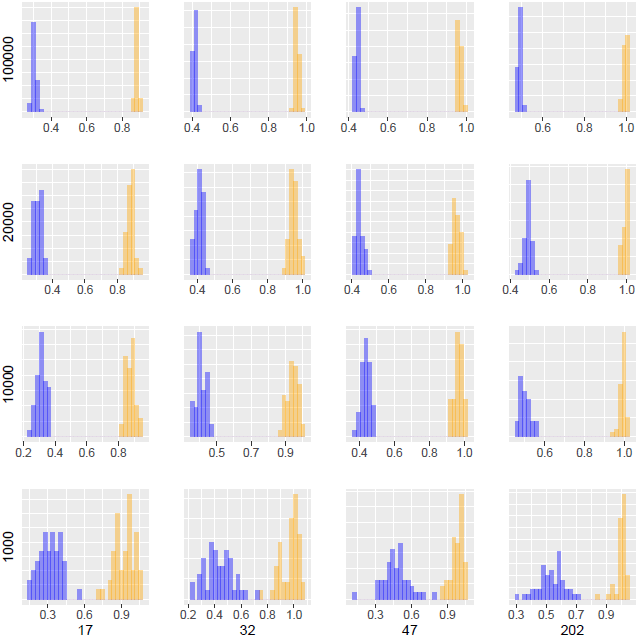
Each panel represents a different combination of *N* (columns) and *L* (rows) and shows a histogram for the estimates of simulated relatedness of *r*_*s*_ = 1.0 and *r*_*s*_ = 0.5 over 50 independent datasets for pedigree *f1.0*.

**Figure SF3:**
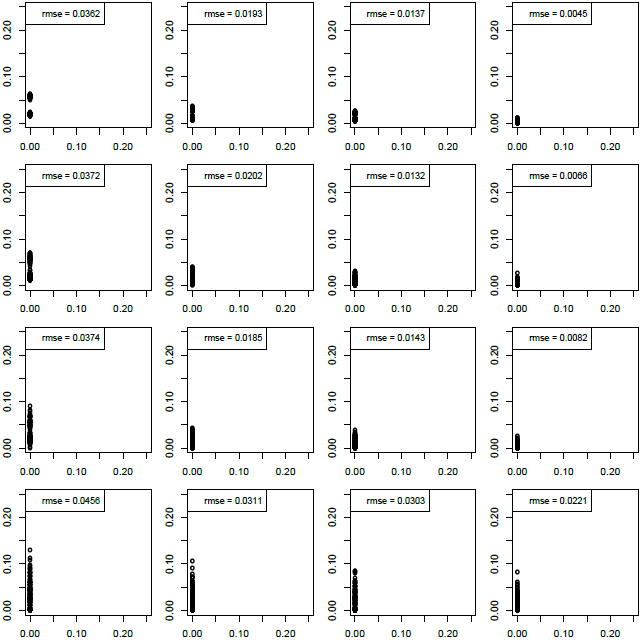
Each panel represents a different combination of *N* (columns) and *L* (rows) and shows an x-y plot for estimated (x axis) and simulated (y axis) contamination rates for pedigree *f1.0*. *C*_*i*_ was simulated to be 0.

**Figure SF4:**
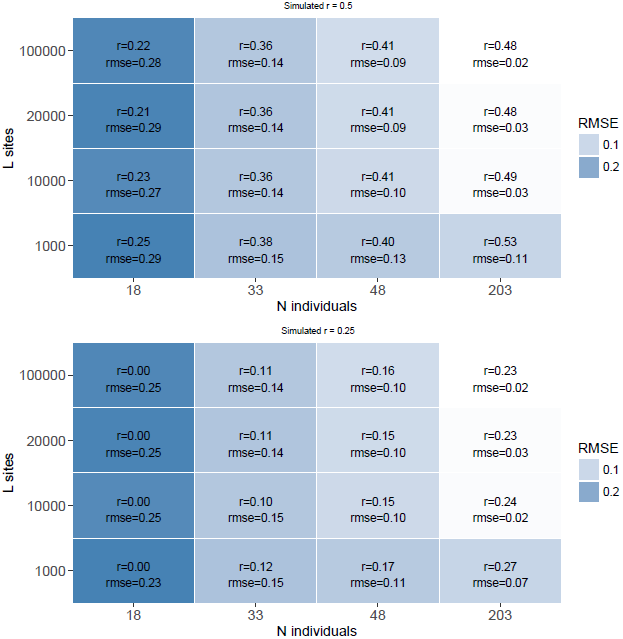
Similar setting as in figure SF1 but for pedigree *f2.0* with *r*_*s*_ = 0.5 (top panel) and *r*_*s*_ = 0.25 (bottom panel). *C*_*i*_ = 0, e = 0.

**Figure SF5:**
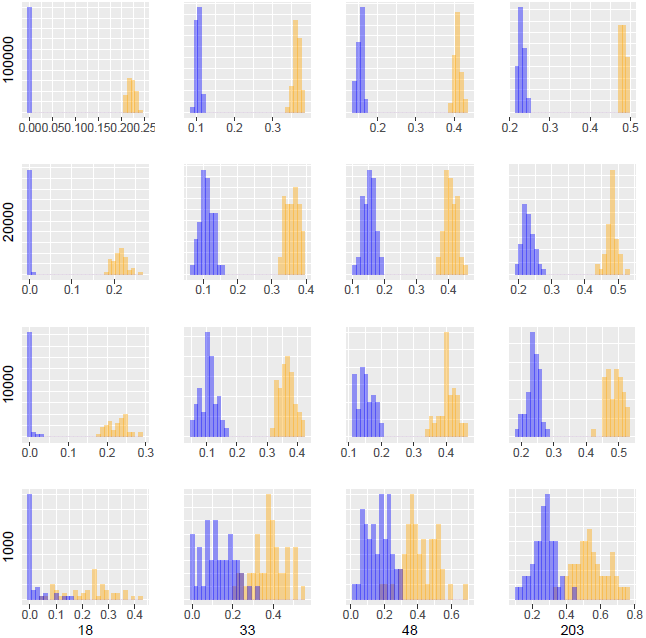
Similar setting as in figure SF2 but for pedigree *f2.0* with *r*_*s*_ = 0.5 and *r*_*s*_ = 0.25. *C*_*i*_ = 0, e = 0.

**Figure SF6:**
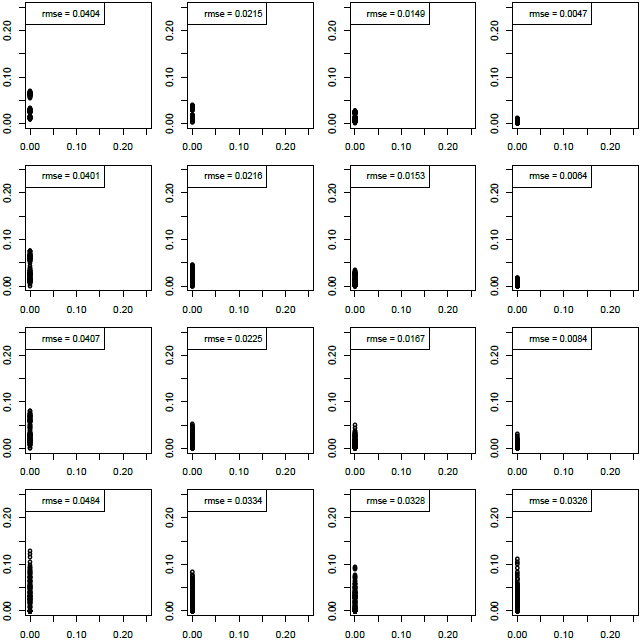
Similar setting as in figure SF3 but for pedigree *f2.0* with *r*_*s*_ = 0.5 and *r*_*s*_ = 0.25. *C*_*i*_ = 0, e = 0.

**Figure SF7:**
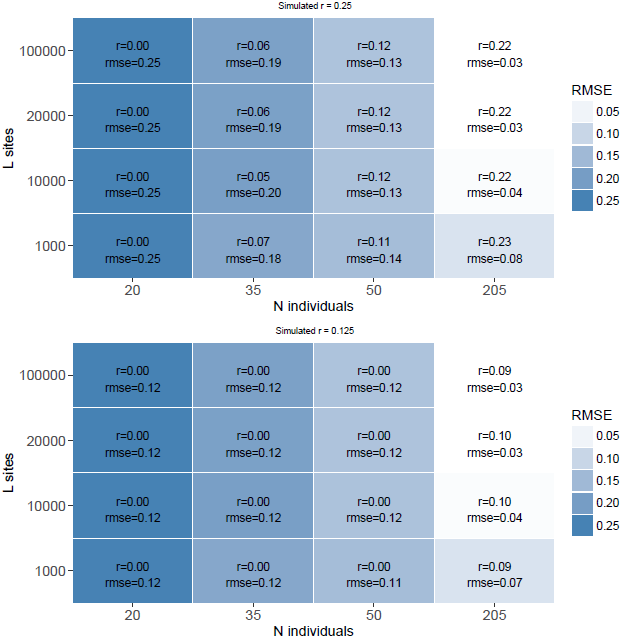
Similar setting as in figure SF1 but for pedigree *f3.0* with *r*_*s*_ = 0.25 (top panel) and *r*_*s*_ = 0.125 (bottom panel). *C*_*i*_ = 0, e = 0.

**Figure SF8:**
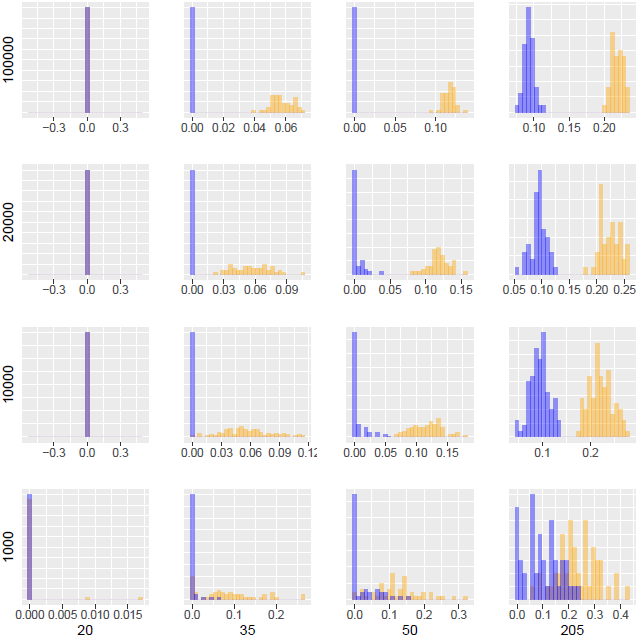
Similar setting as in figure SF2 but for pedigree *f3.0* with *r*_*s*_ = 0.25 and *r*_*s*_ = 0.125. *C*_*i*_ = 0, e = 0.

**Figure SF9:**
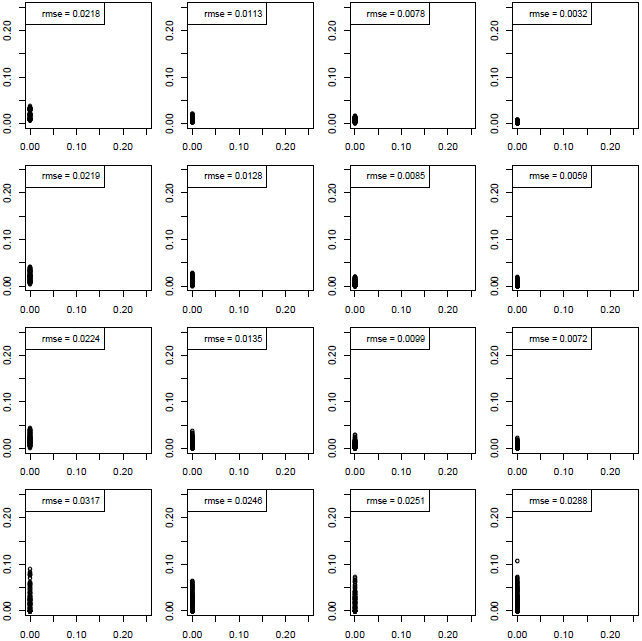
Similar setting as in figure SF3 but for pedigree *f3.0* with *r*_*s*_ = 0.25 and *r*_*s*_ = 0.125. *C*_*i*_ = 0, e = 0.

**Figure SF10:**
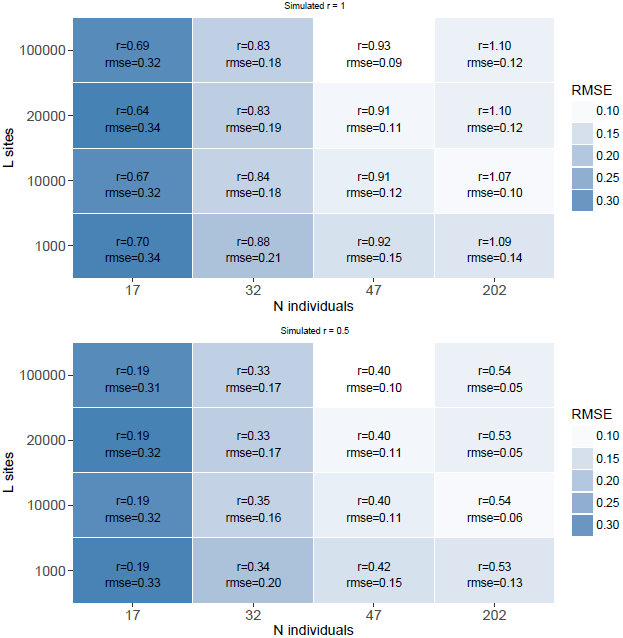
Similar setting as in figure SF1 but *C*_*i*_ is between 2% and 25%, e = 0.001.

**Figure SF11:**
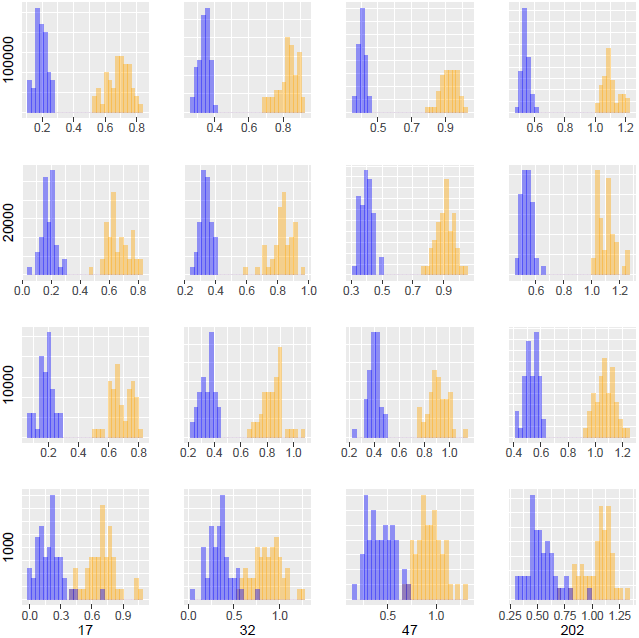
Similar setting as in figure SF3 but *C*_*i*_ is between 2% and 25%, e = 0.001.

**Figure SF12:**
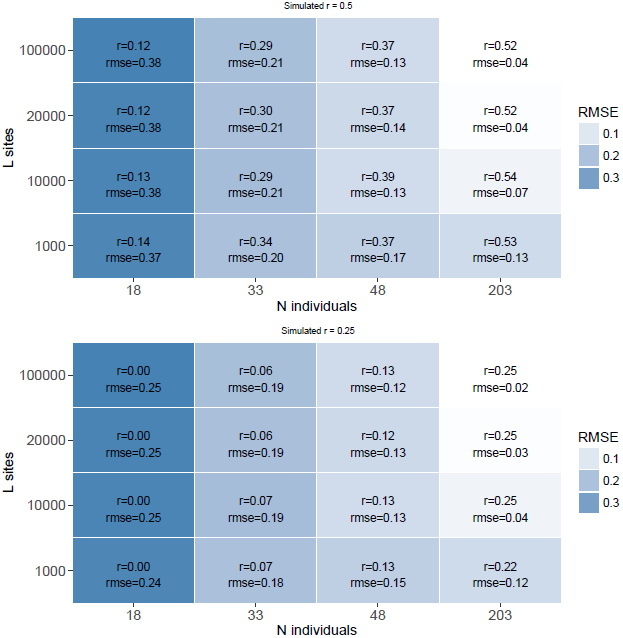
Similar setting as in figure SF4 but *C*_*i*_ is between 2% and 25%, e = 0.001.

**Figure SF13:**
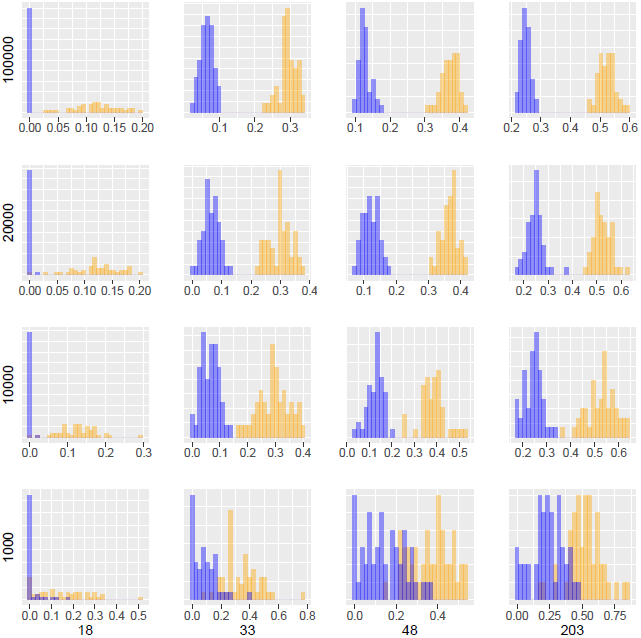
Similar setting as in figure SF5 but *C*_*i*_ is between 2% and 25%, e = 0.001.

**Figure SF14:**
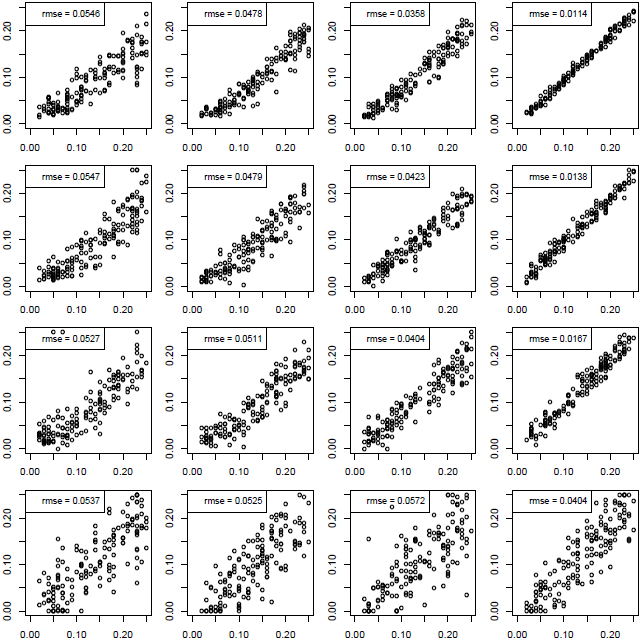
Similar setting as in figure SF6 but *C*_*i*_ is between 2% and 25%, e = 0.001.

**Figure SF15:**
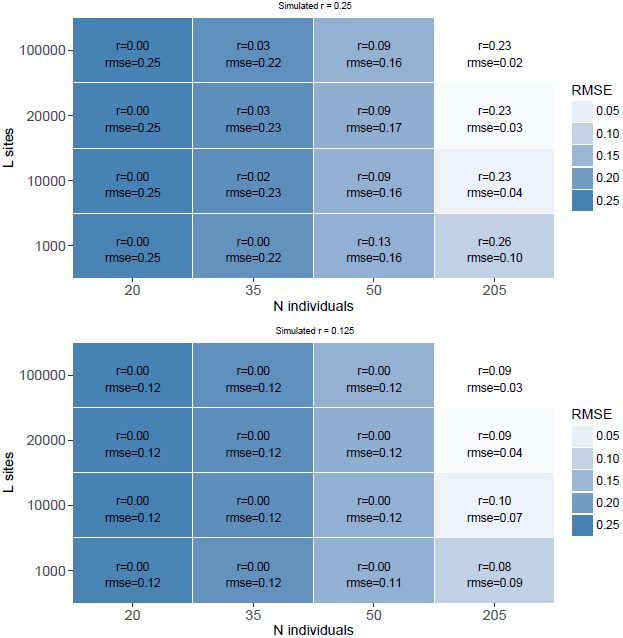
Similar setting as in figure SF7 but *C*_*i*_ is between 2% and 25%, e = 0.001.

**Figure SF16:**
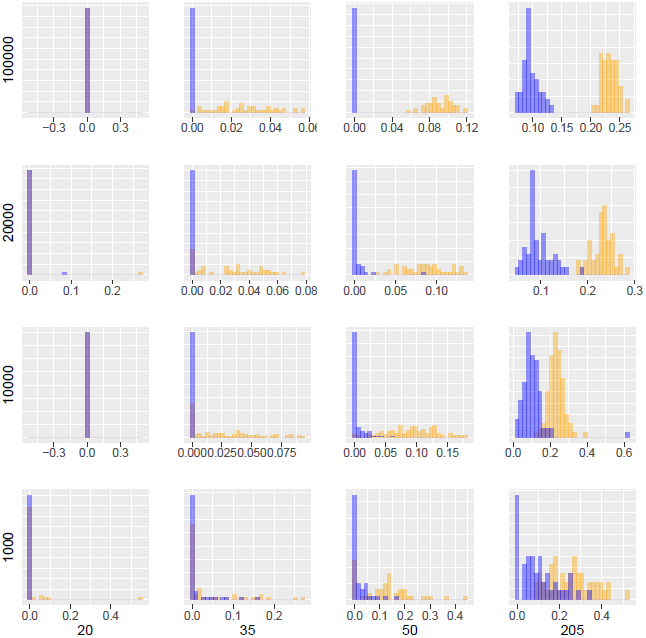
Similar setting as in figure SF8 but *C*_*i*_ is between 2% and 25%, e = 0.001.

**Figure SF17:**
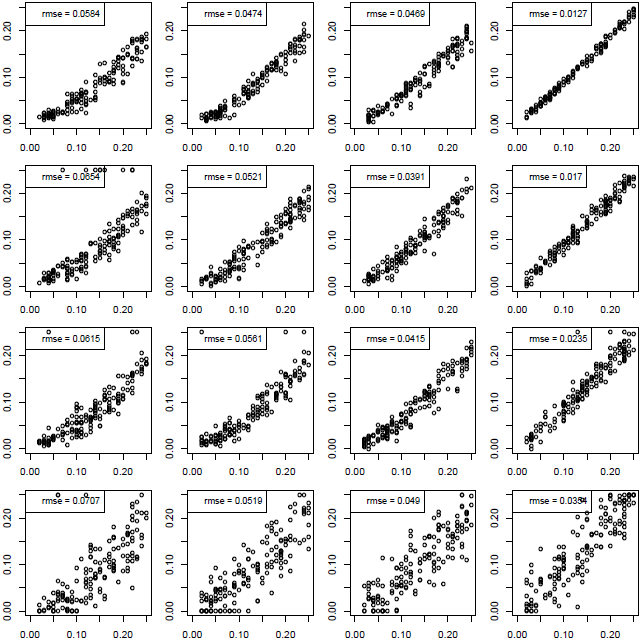
Similar setting as in figure SF9 but *C*_*i*_ is between 2% and 25%, e = 0.001.

**Figure SF18:**
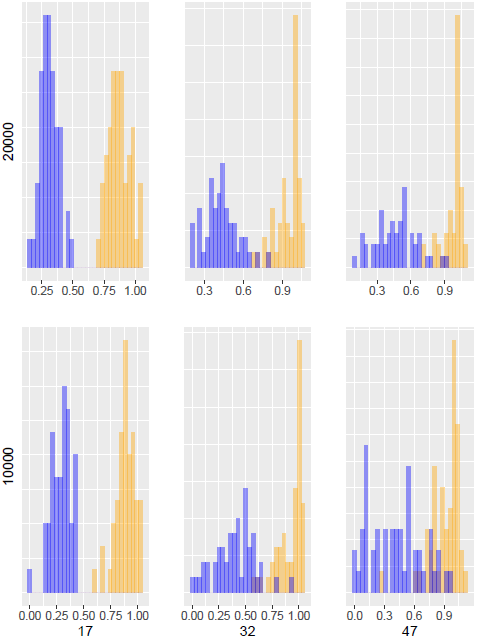
Each panel represents a different combination of N (columns) and L (rows) and shows a histogram for the estimates of simulated relatedness of rs = 1.0 and rs = 0.5 over 50 independent datasets for pedigree f1.0. However, at each genomic site read information was available only from a random subset of 5 different individuals (out of a total of N).

**Figure SF19:**
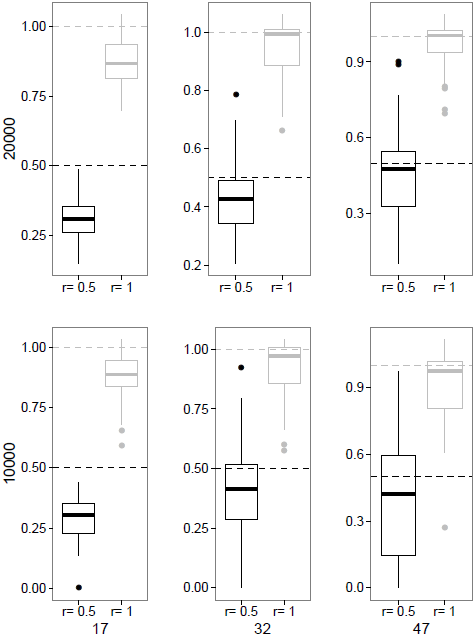
Each panel represents a different combination of N (columns) and L (rows) and shows a boxplot for the estimates of simulated relatedness of rs = 1.0 and rs = 0.5 over 50 independent datasets for pedigree f1.0. Dashed horizontal lines denote the simulated values of rs. However, at each genomic site read information was available only from a random subset of 5 different individuals (out of a total of N).

**Figure SF20:**
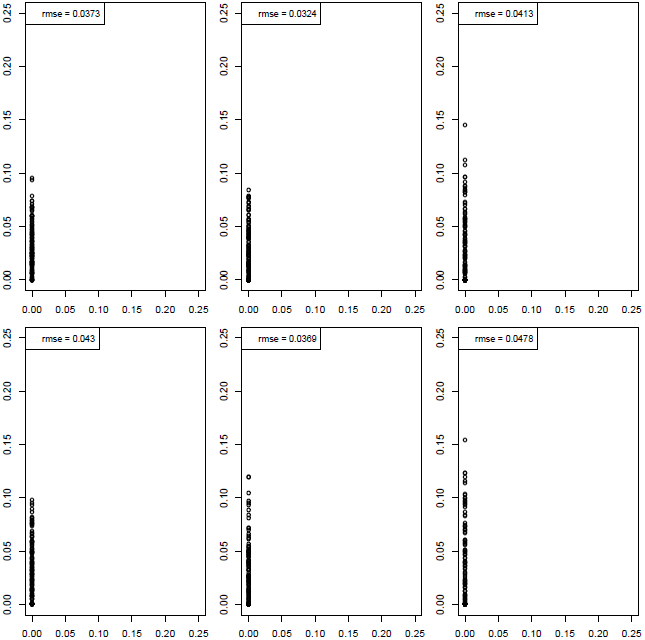
Each panel represents a different combination of N (columns) and L (rows) and shows an x-y plot for estimated (x axis) and simulated (y axis) contamination rates for pedigree f1.0. Ci was simulated to be 0. However, at each genomic site read information was available only from a random subset of 5 different individuals (out of a total of N).

**Figure SF21:**
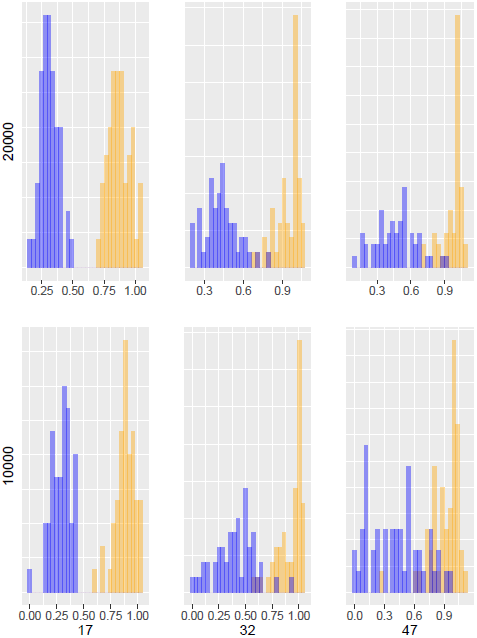
Similar setting as in figure SF18 but for pedigree *1.1*.

**Figure SF22:**
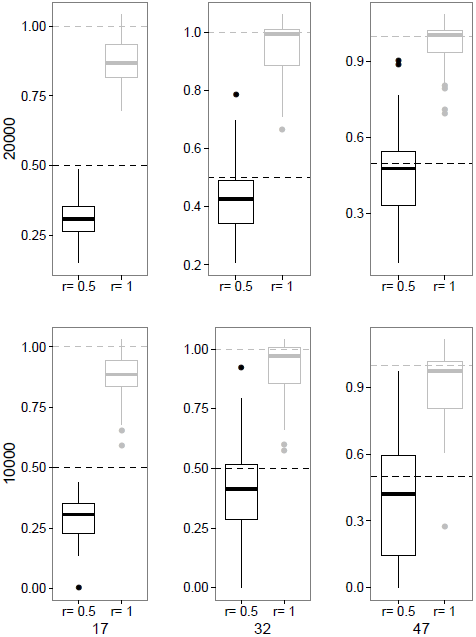
Similar setting as in figure SF19 but for pedigree *1.1*.

**Figure SF23:**
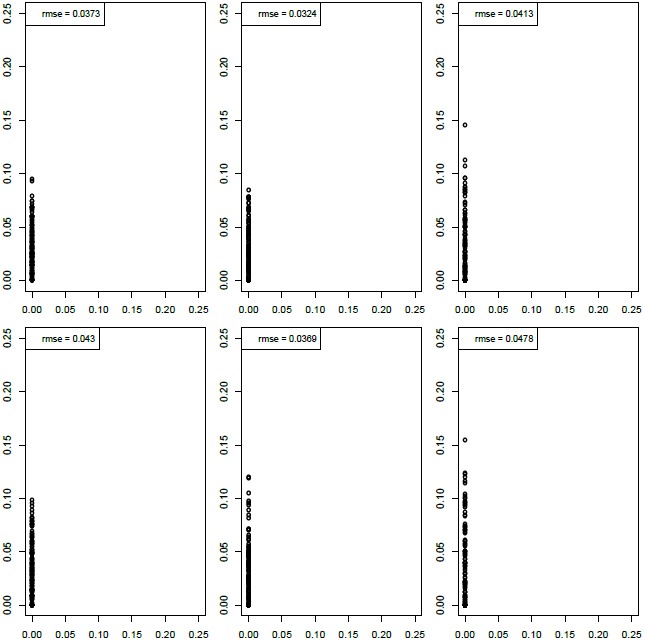
Similar setting as in figure SF20 but for pedigree *1.1*.

**Figure SF24:**
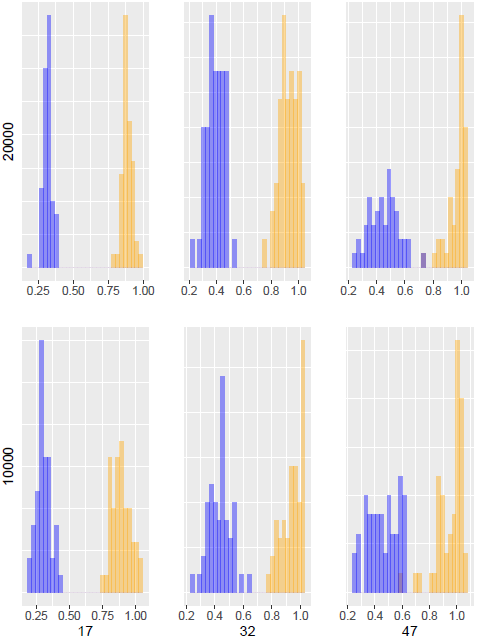
Each panel represents a different combination of N (columns) and L (rows) and shows a histogram for the estimates of simulated relatedness of rs = 1.0 and rs = 0.5 over 50 independent datasets for pedigree f1.0. However, at each genomic site read information was available only from a random subset of 10 different individuals (out of a total of N).

**Figure SF25:**
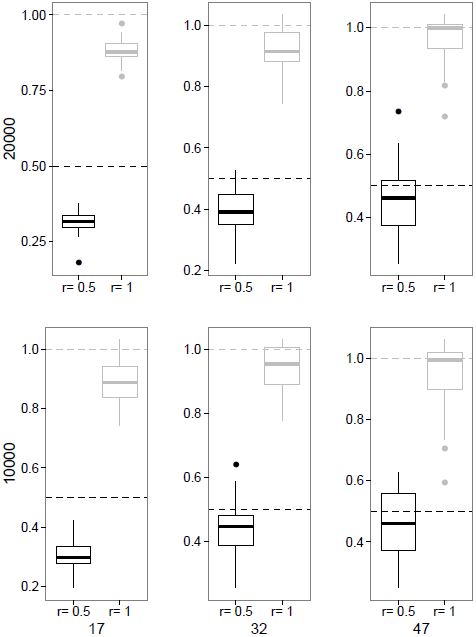
Each panel represents a different combination of N (columns) and L (rows) and shows a boxplot for the estimates of simulated relatedness of rs = 1.0 and rs = 0.5 over 50 independent datasets for pedigree f1.0. Dashed horizontal lines denote the simulated values of rs. However, at each genomic site read information was available only from a random subset of 10 different individuals (out of a total of N).

**Figure SF26:**
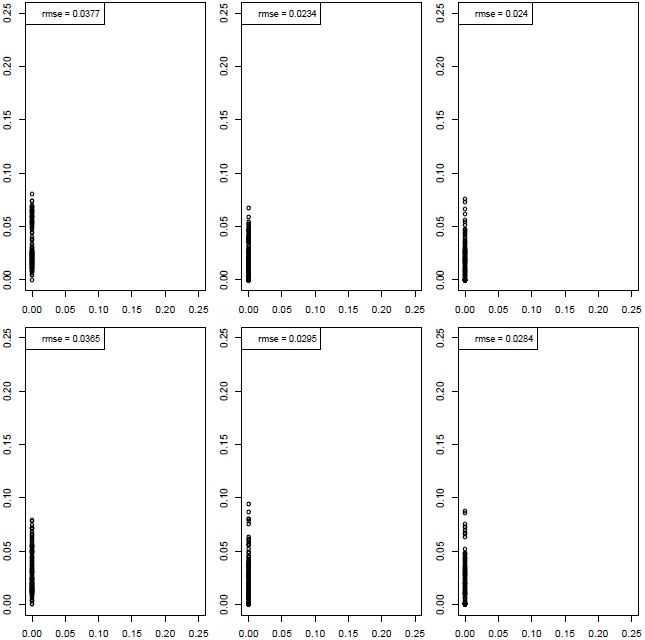
Each panel represents a different combination of N (columns) and L (rows) and shows an x-y plot for estimated (x axis) and simulated (y axis) contamination rates for pedigree f1.0. Ci was simulated to be 0. However, at each genomic site read information was available only from a random subset of 10 different individuals (out of a total of N).

**Figure SF27:**
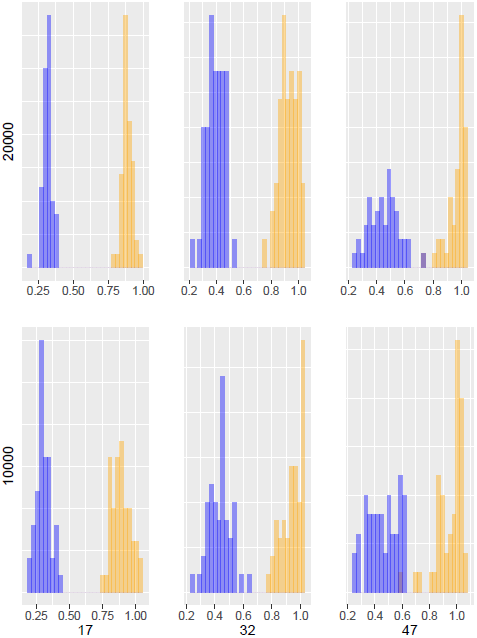
Similar setting as in figure SF24 but for pedigree *1.1*.

**Figure SF28:**
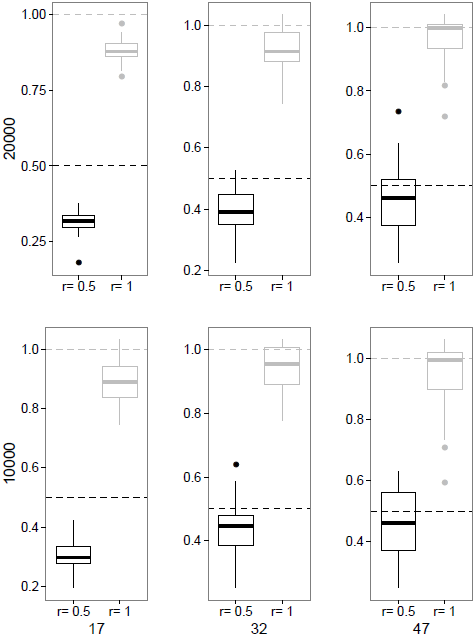
Similar setting as in figure SF25 but for pedigree *1.1*.

**Figure SF29:**
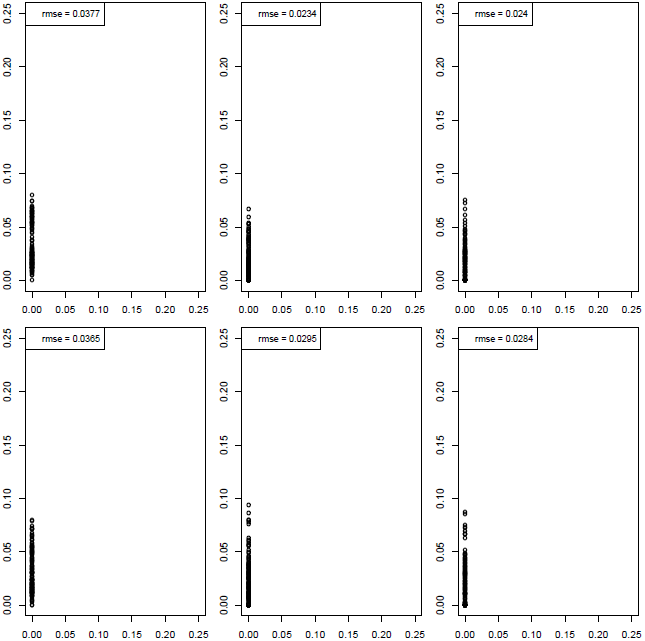
Similar setting as in figure SF26 but for pedigree *1.1*.

**Figure SF30:**
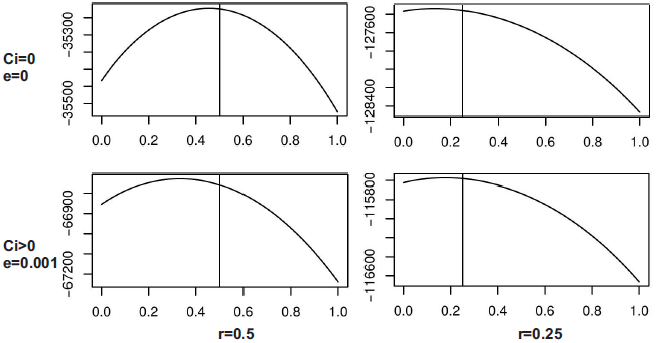
This figure shows conditional likelihoods for results based on *r*_*s*_ = 0.25 and *r*_*s*_ = 0.5 under the presence and absence of contamination rate and sequencing error.

